# An analytical solution simulating growth of Lewy bodies

**DOI:** 10.1101/2021.12.22.473938

**Authors:** Ivan A. Kuznetsov, Andrey V. Kuznetsov

## Abstract

This paper reports a minimal model simulating the growth of a Lewy body (LB). To the best of our knowledge, this is the first model simulating Lewy body growth. The LB is assumed to consist of a central spherical core, which is composed of membrane fragments and various dysfunctional intracellular organelles, and a halo, which is composed of alpha-synuclein fibrils. Membrane fragments and alpha-synuclein monomers are assumed to be produced in the soma at constant rates. The growth of the core and the halo are simulated by the Finke-Watzky model. Analytical (closed-form) solutions describing the growth of the core and the halo are obtained. A sensitivity analysis in terms of model parameters is performed.

## 1. Introduction

The formation of Lewy bodies (LBs) in the soma of dopaminergic neurons is one of the main characteristic features of Parkinson’s disease (PD) (Kalia & Lang, 2015; Gomez-Benito *et al.*, 2020; Mahul-Mellier *et al.*, 2020). Until recently it was believed that LBs mostly consist of misfolded forms of alpha-synuclein (α-syn) (Spillantini *et al.*, 1998). However, new data reported in Shahmoradian *et al.* (2019) show that LBs are mostly composed of various lipid components, membrane fragments, organelles, vesicular structures, dysmorphic mitochondria, autophagosome precursors, and lysosomes. The reasons for this surprising observation and the question of whether α-syn fibrils are a part of LBs are heavily debated (Lashuel, 2020; Fares *et al.*, 2021).

In this paper, we develop a model that can help shed light on these previous observations. It is known that brainstem-type LBs consist of a dense core surrounded by a halo composed of radiating filaments (Mahul-Mellier *et al.*, 2020; Fares *et al.*, 2021). The halo resembles a spherical shell. LBs are likely formed from their precursors, pale bodies, which are the granular structures that lack a halo (Fares *et al.*, 2021). A pale body can progress to become a classical LB with a halo (Shults, 2006).

It is still debated whether LBs are neurotoxic or neuroprotective. A large body of research argues that LBs are cytotoxic (Power *et al.*, 2017). On the other hand, many researchers support the view that LBs remove cytotoxic or unwanted species from the cytosol and thus are neuroprotective (Olanow *et al.*, 2004). Parkkinen *et al.* (2011) reported that in PD the loss of neurons in the ventrolateral tier of the pars compacta of the substantia nigra (SN) is much higher than the number of LBs. LB-containing neurons seem to be morphologically healthier than neighboring non-LB-containing neurons. It seems that most SN neurons that die of apoptosis do not contain LBs. On the other hand, Shults (2006) noted that if LBs are neuroprotective, it is hard to explain why as PD progresses the number of neurons with LBs in SN pars compacta does not increase. Instead, LB-laden neurons die.

To address this paradox, in our model we assumed that the process of formation of LBs includes two stages. First, a structure composed of membranous fragments, organelles, vesicular structures, and various lipid constituents is formed by aggregation of these components into the core of an LB. We assumed that the core is benign to the neuron. Then, surface-initiated polymerization (Gambinossi *et al.*, 2015) induced by the surface of the core leads to the growth of α-syn fibrils that form the halo of the LB (Fig. 1). α-syn fibrils elongate by adding α-syn monomers from the cytosol. We assumed that radiating α-syn filaments are cytotoxic. These radiating filaments also prevent further growth of the core by cutting off the supply of membrane fragments to the core, thus blocking broken intracellular material from being removed from the cytosol by incorporating it into the core of the LB.

**Fig. 1.**
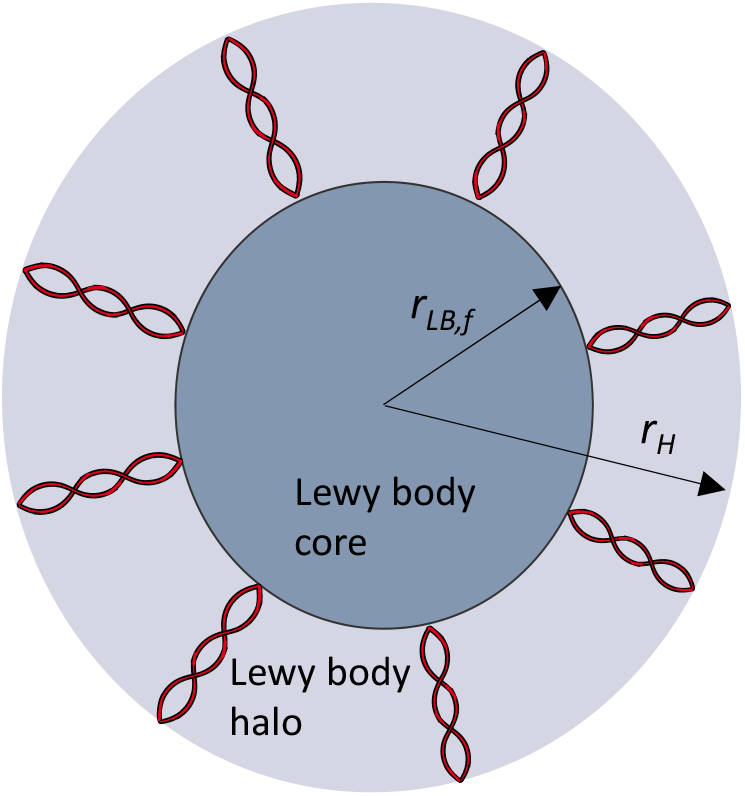
A diagram showing an LB composed of two regions: the central core composed of dense granular material and the halo containing radiating filaments (modeled after an electron micrograph shown in Fig. 1 in Olanow *et al.* (2004) and Fig. 3 in Fares *et al.* (2021)) *Figure generated with the aid of servier medical art, licensed under a creative common attribution 3.0 generic license. http://Smart.servier.com.*

In this paper, we develop a mathematical model based on these assumptions and obtain its analytical solution.

## 2. Materials and models

### 2.1 Governing equations

Independent variables in the model are summarized in Table 1, dependent variables in the model are summarized in Table 2, and parameters involved in the model are summarized in Table 3.

**Table 1.** Independent variables in the model.

| Symbol | Definition | Units |
| --- | --- | --- |
| $T$ | Time | s |

**Table 2.** Dependent variables in the model.

| Symbol | Definition | Units |
| --- | --- | --- |
| $[A_{FR}]$ | Molar concentration of lipid membrane fragments (as well as damaged organelles, lysosomes, and damaged mitochondria; generally, any fragments that can form the core of the LB) in the soma | $\mu\text{M}$ |
| $[B_{FR}]$ | Molar concentration of aggregated lipid membrane fragments in the core of the LB | $\mu\text{M}$ |
| $[A_{AS}]$ | Molar concentration of α-syn monomers in the soma | $\mu\text{M}$ |
| $[B_{AS}]$ | Molar concentration of α-syn fibrils in the halo of the LB | $\mu\text{M}$ |

**Table 3.**
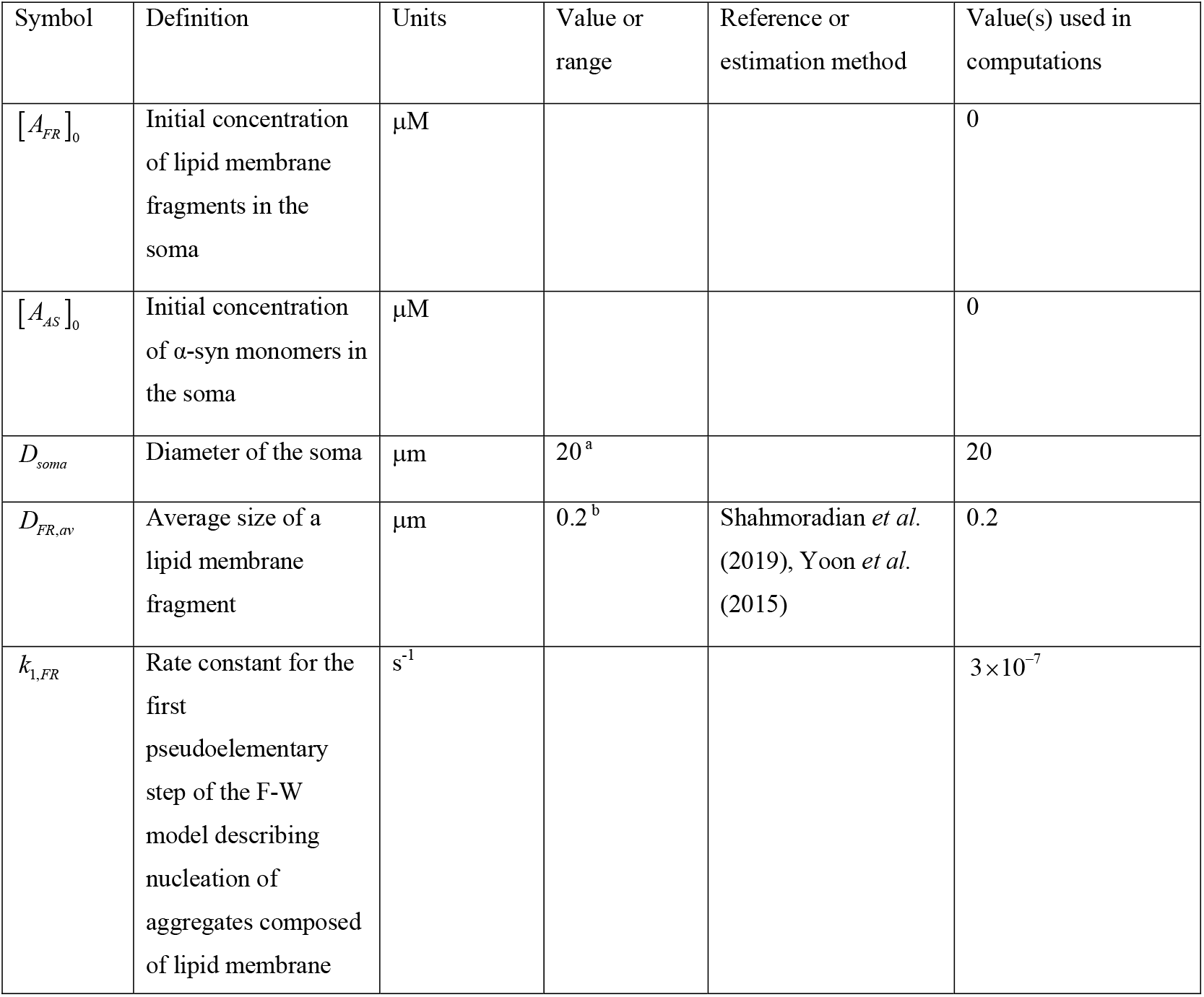
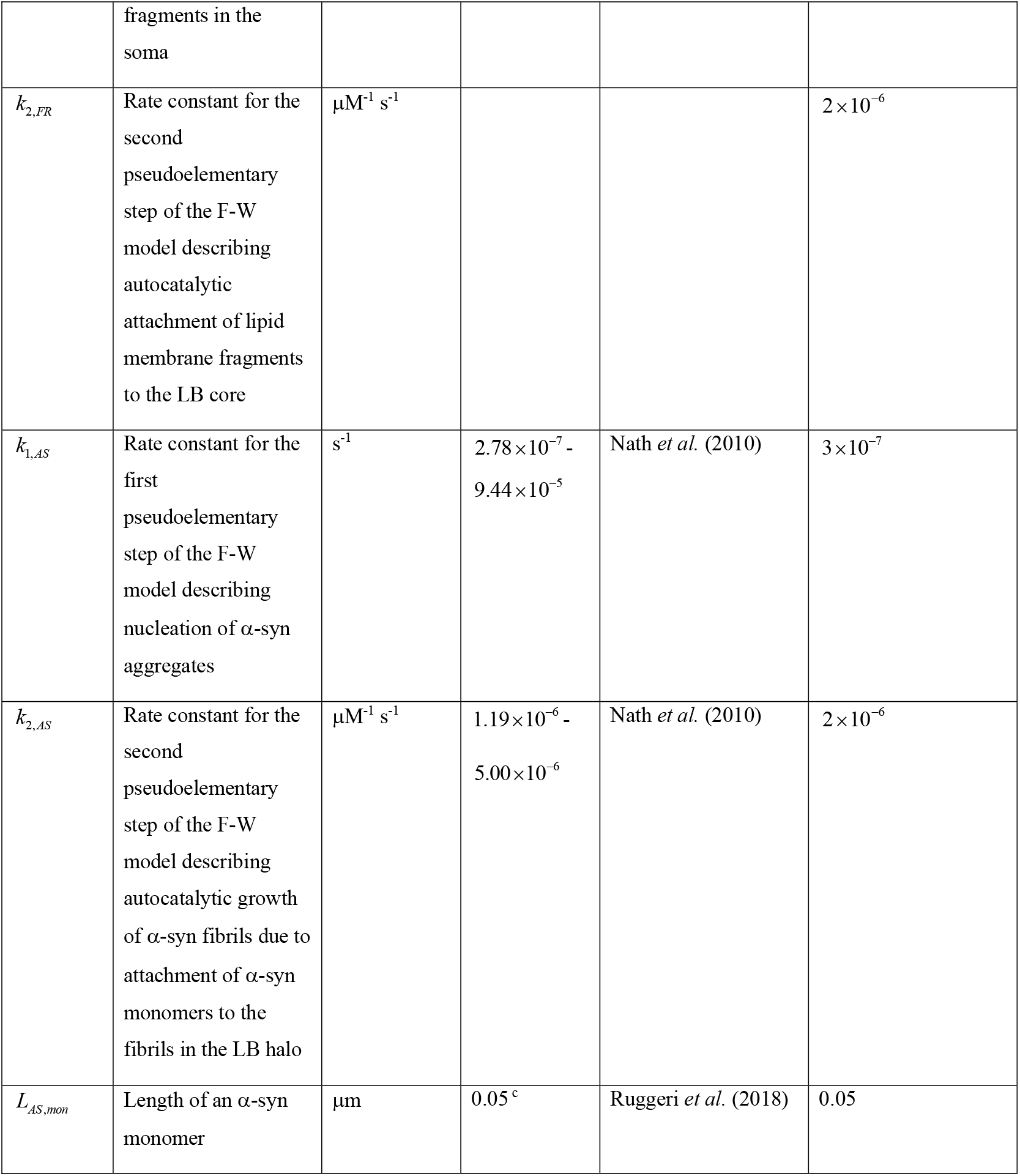
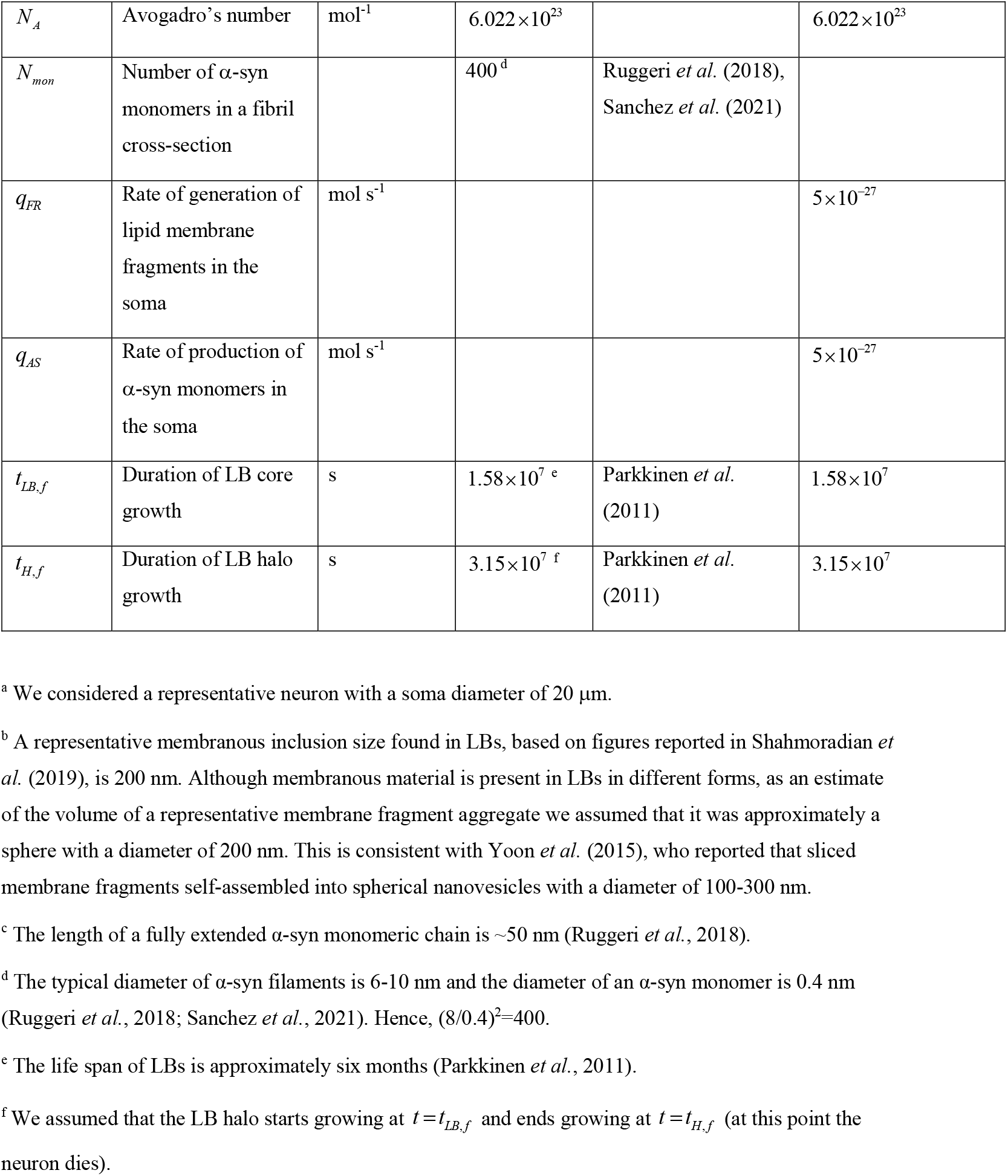
Parameters involved in the model.

Since the formation of LBs is a very slow process, we assumed that it is limited by the kinetics of two aggregation processes: first, by the kinetics of membrane fragment aggregation into the LB core and then by the kinetics of α-syn fibril growth that forms the LB halo.

A minimalistic 2-step Finke-Watzky (F-W) model describes continuous nucleation followed by fast, autocatalytic surface growth. This model is described by the following transitions:

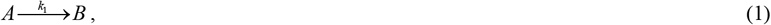

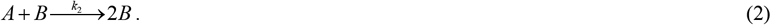

Usually, the F-W model is applied to simulate the conversion of a given initial concentration of monomers, [*A*]_0_, into polymers, [*B*]. Analytical solutions for such cases are reported, for example, in Bentea *et al.* (2017). Here we apply the F-W model to a different situation. Results reported in Shahmoradian *et al.* (2019) and the analysis reported in Fares *et al.* (2021) suggest that the core of the LB is composed of lipid membrane fragments, damaged organelles, lysosomes, and damaged mitochondria. Hereafter, for briefness, we will use the term membrane fragments to denote any fragmented/damaged components that can form the core of an LB.

We assume that membrane fragments are produced at a rate *q_FR_* in the soma. Stating the conservation of membrane fragments in the soma results in the following equation:

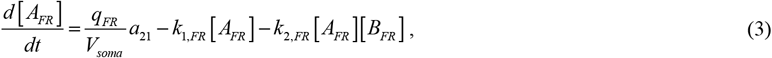

where 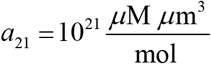 is the conversion factor from 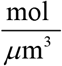 to *μ*M. The first term on the right-hand side of Eq. (3) describes the rate of production of membrane fragments, while the second term describes the rate of conversion of membrane fragments into aggregates by nucleation. The third term describes the rate of conversion of membrane fragments into aggregates by autocatalytic growth. The second and third terms on the right-hand side of Eq. (3) are written by applying the law of mass action to express the rates of reactions given by Eqs. (1) and (2), respectively.

Stating the conservation of membrane aggregates in the soma, which form the core of the LB, leads to the following equation:

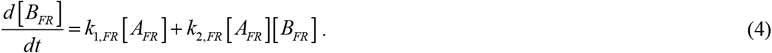

On the right-hand side of Eq. (4), the first term describes the rate of production of aggregates by nucleation and the second term describes the rate of production of aggregates by autocatalytic growth. These terms have the same magnitude but opposite signs as the second and third terms on the right-hand side of Eq. (3). This is because in the F-W model the rate of production of aggregates is equal to the rate of monomer disappearance.

Eqs. (3) and (4) are solved subject to the following initial condition:

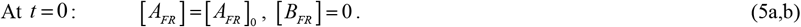

By adding Eqs. (3) and (4) and integrating the result with respect to time, the following is obtained:

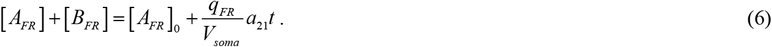

The increase of [*A_FR_*] + [*B_FR_*] with time is due to the production of membrane fragments in the soma. Eliminating [*A_FR_*] from Eq. (4) by using Eq. (6), the following is obtained:

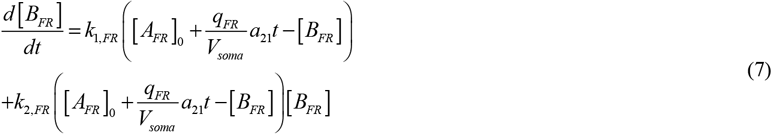

The solution of Eq. (7) subject to initial condition (5b) was obtained using the DSolve function followed by the FullSimplify function in Mathematica 13.3 (Wolfram Research, Champaign, IL). The obtained solution is

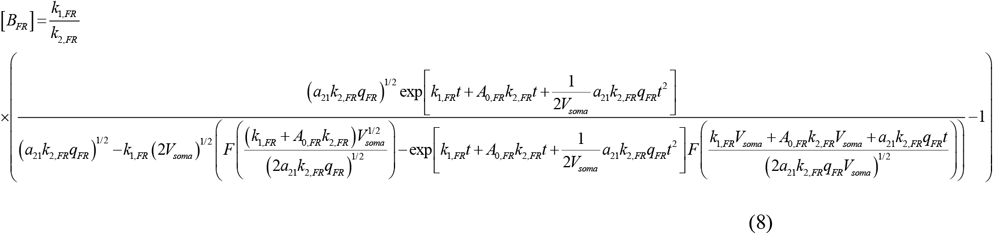

Here *F*(*x*) is Dawson’s integral:

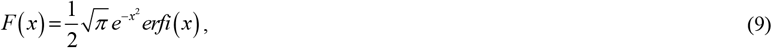

where *erfi*(*x*) is the imaginary error function.

The growth of the core of an LB (Fig. 1) is simulated as follows. The volume of the core of the LB can be found as:

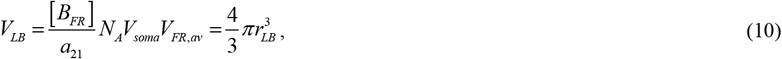

where

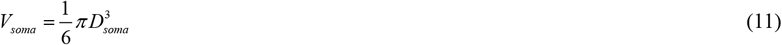

is the volume of the soma and

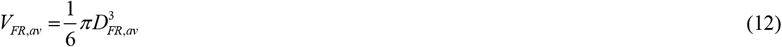

is the average volume of a lipid membrane fragment.

This is consistent with (Szabo & Lente, 2019), which pointed out that the volume of a nanoparticle is proportional to the number of monomers making up the particle. Hence, the size of a nanoparticle is proportional to the cubic root of the number of monomers composing the particle.

The growth continues until *r_LB_* reaches *r_LB,f_*, where

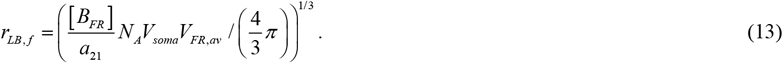

We assumed that α-syn monomers are produced at a rate *q_AS_* in the soma. We also assumed that α-syn fibrils are produced by the polymerization of α-syn monomers in the halo region of the LB. The F-W model given by Eqs. (1) and (2) is utilized again. Stating the conservation of α-syn monomers in the soma results in the following equation:

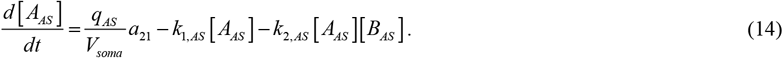

Stating the conservation of α-syn fibrils, which form the halo region of the LB, leads to the following equation:

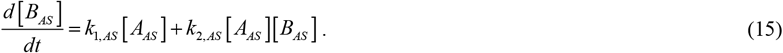

Eqs. (14) and (15) are written similarly to Eqs. (3) and (4). The first term of the right-hand side of Eq. (14) describes the rate of production of α-syn monomers. The second and third terms simulate the rates of conversion of α-syn monomers into fibrils by nucleation and autocatalytic growth, respectively. The two terms on the right-hand side of Eq. (15) simulate the rate of increase of α-syn fibril concentration due to nucleation and autocatalytic growth, respectively.

Eqs. (14) and (15) are solved subject to the following initial condition:

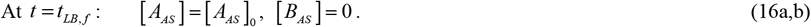

The initial value problem described by Eqs. (14)-(16) becomes similar to that described by Eqs. (3)-(5) if the transformation

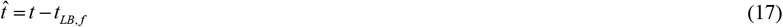

is utilized.

By adding Eqs. (14) and (15) and integrating the result with respect to time, the following is obtained:

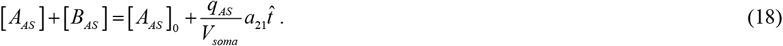

The increase of [*A_AS_*] + [*B_AS_*] with time is due to the production of membrane fragments in the soma. Eliminating [*A_AS_*] from Eq. (15) by using Eq. (18), the following is obtained:

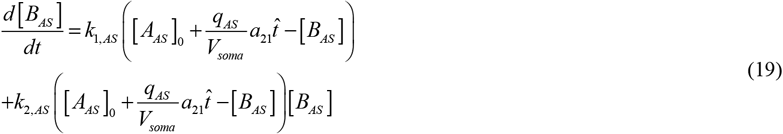

The solution of Eq. (19) subject to initial condition (16b) is

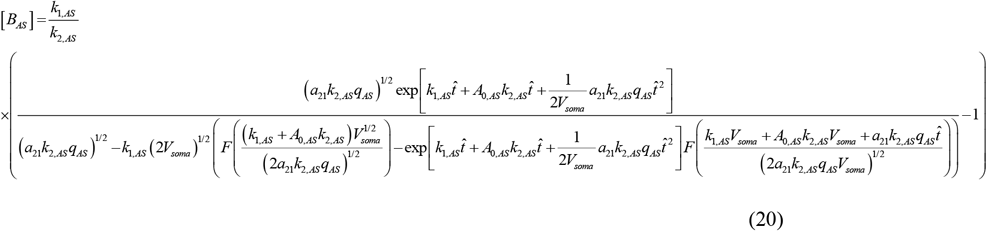

The thickness of the halo of the LB (Fig. 1) in the radial direction can then be found as:

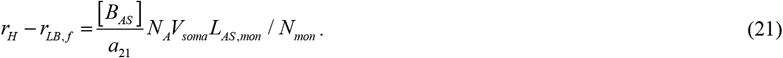

### 2.2 Sensitivity analysis

We investigated how the outer halo radius at the end of the simulation (*t_H,f_* = 1 year) depends on various model parameters. To perform this analysis, we computed the local sensitivity coefficients, which are first-order partial derivatives of the outer halo radius with respect to the parameters (Beck & Arnold, 1977; Zadeh & Montas, 2010; Zi, 2011; Kuznetsov & Kuznetsov, 2019). The sensitivity coefficient of *r_H,f_* to parameter *q_FR_*, for example, was calculated as follows:

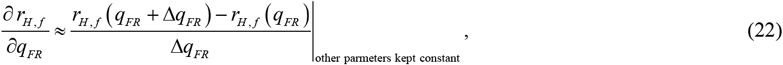

where Δ*q_FR_* = 10^-3^ *q_FR_* is the step size. To test the independence of the sensitivity coefficients to the step size we tested various step sizes.

Non-dimensionalized relative sensitivity coefficients were calculated following Zadeh & Montas (2010), Kacser *et al.* (1995) as, for example:

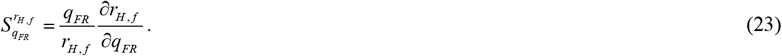

## 3. Results

### 3.1. Analysis of the model

The increase of the generation rate of lipid membrane fragments, FR *q_FR_*, leads to an increase in the concentration of these fragments in the soma (Fig. 2a) and faster growth of the core of the LB (Fig. 2b). After six months of growth, the LB lipid core reaches a diameter that is in the range 5.02 μm - 8.03 μm, depending on the value of *q_FR_*. After another six months (the total time of LB growth is assumed to be one year, after that the neuron dies), the outer diameter of the halo reaches 9.80 μm-12.8 μm, which is consistent with the 8-30 μm range of LB diameter reported in Olanow et al. (2004).

**Fig. 2.**
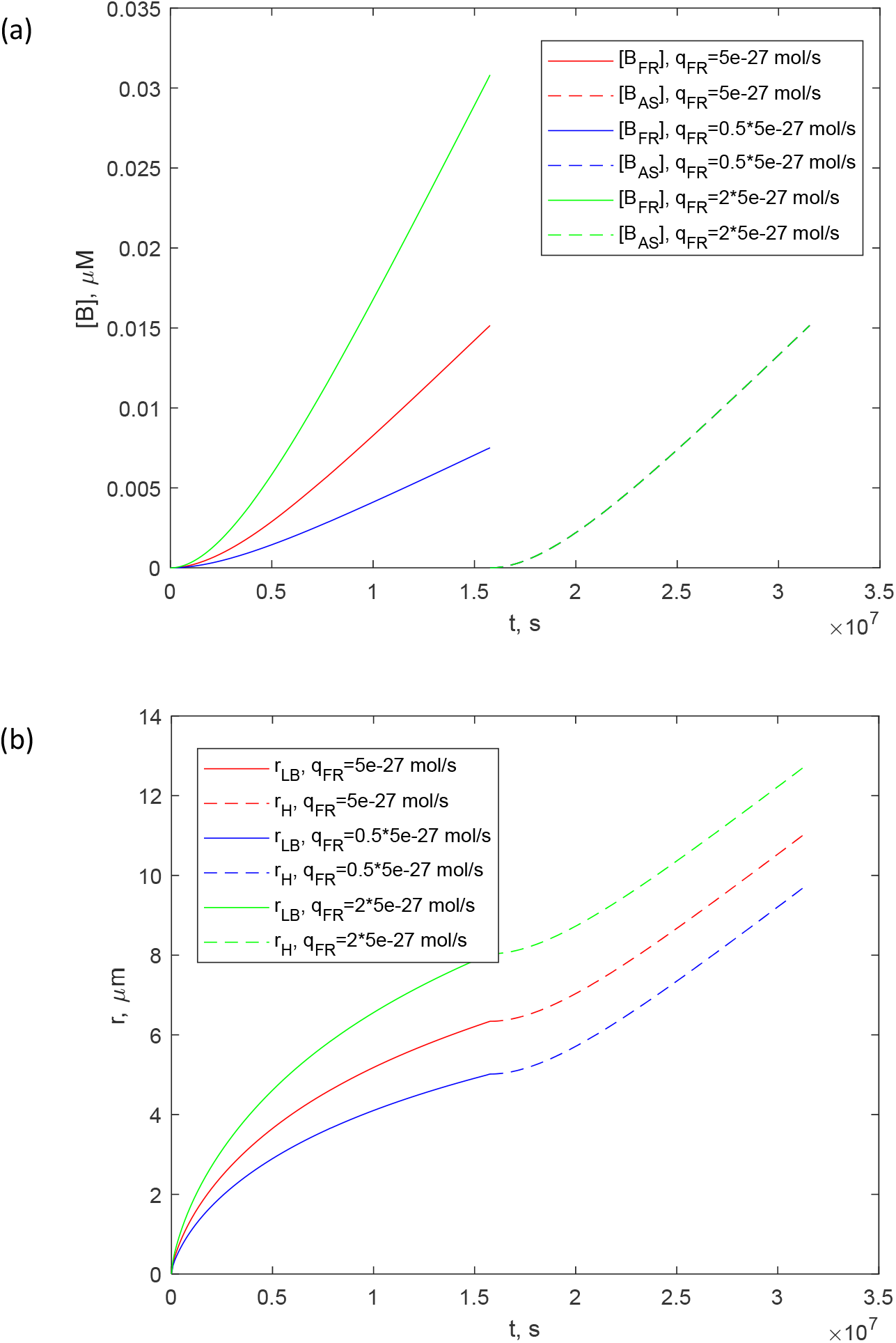
*(a)* Molar concentration of aggregated lipid membrane fragments in the core of the LB, [*B_FR_*], and molar concentration of α-syn fibrils in the halo of an LB,[*B_AS_*], versus time. (b) Radius of the core of the LB, *r_LB_*, and the outer radius of the halo of the LB versus time, *r_H_*, for three values of *q_FR_*. *q_AS_* = 5 × 10^-27^ mol s^-1^, *k*_1,*FR*_ = 3×10^-7^ s^-1^, *k*_2,*FR*_ = 2×10^-6^ μM^-1^ s^-1^, *k*_1,*AS*_ = 3×10^-7^ s^-1^, *k*_2,*AS*_ = 2×10^-6^ μM^-1^ s^-1^. The MATLAB notation is utilized, i.e., 5e-27 means 5 × 10^-27^.

An increase in the rate of production of α-syn monomers in the soma, *q_AS_*, leads to an increase in the concentration of α-syn monomers in the soma (Fig. 3a) and faster growth of the LB halo (Fig. 3b). After six months of growth, the diameter of the LB core reaches 6.34 μm. After another six months, the outer diameter of the halo reaches 8.71 μm-16.1 μm, depending on the value of *q_AS_*.

**Fig. 3.**
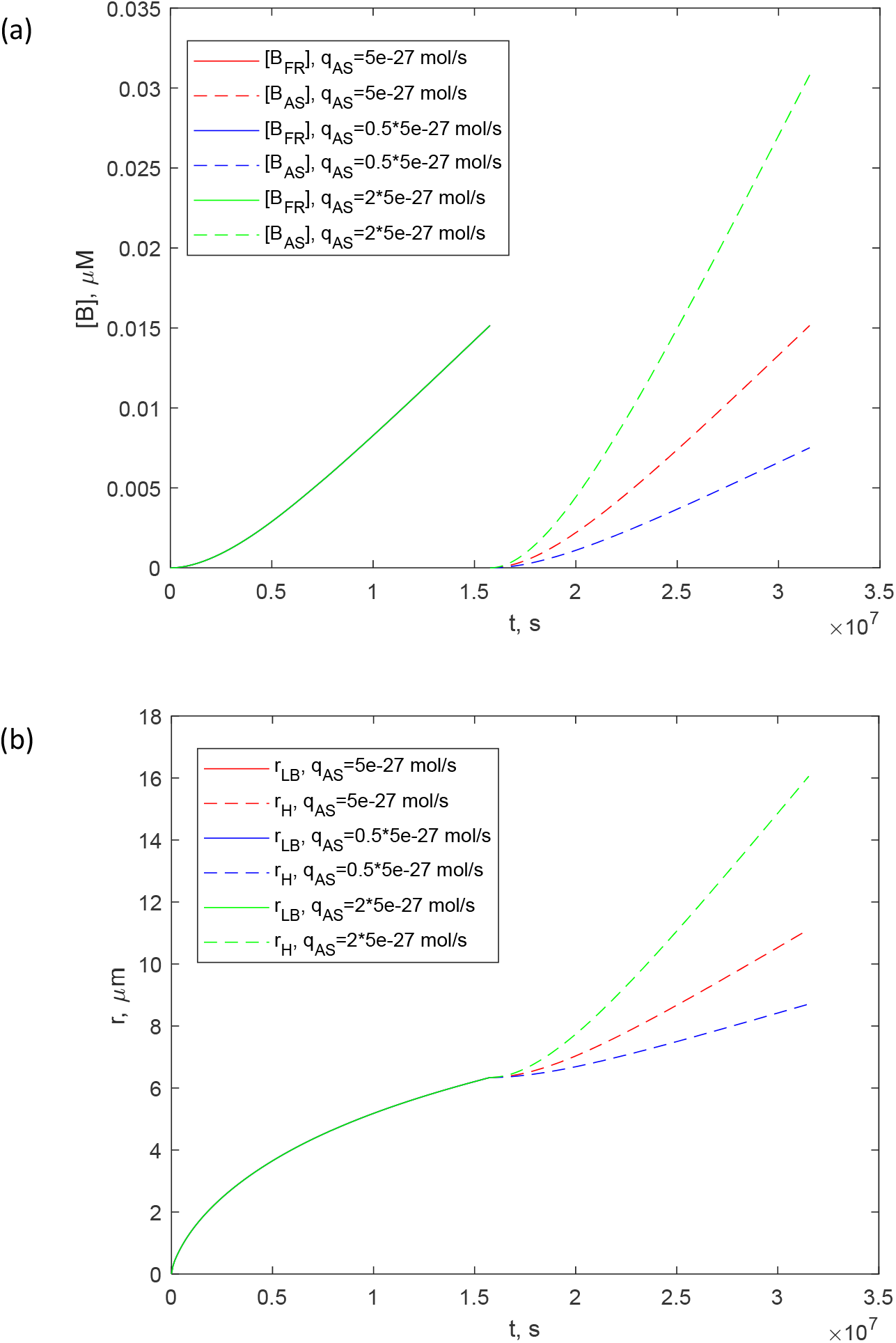
(a) Molar concentration of aggregated lipid membrane fragments in the core of the LB, [*B_FR_*], and molar concentration of α-syn fibrils in the halo of the LB, [*B_AS_*], versus time. (b) Radius of the core of the LB, *r_LB_*, and the outer radius of the halo of the LB versus time, *r_H_*, for three values of *q_AS_*. *q_FR_* = 5 × 10 mol s^-1^, *k*_1,*FR*_ = 3 × 10 s^-1^, *k*_2,*FR*_ = 2 × 10^-6^ μM^-1^ s^-1^, *k*_1,*AS*_ = 3 × 10 s^-1^, *k*_2,*AS*_ = 2 × 10^-6^ μM^-1^ s^-1^.

An increase in the rate constant for the first pseudoelementary step of the F-W model describing nucleation of aggregates composing lipid membrane fragments in the soma, *k*_1,*FR*_, results in faster nucleation of such aggregates (Fig. 4a). This leads to faster growth of the LB core (Fig. 4b). After six months of growth, the diameter of the LB core reaches 5.89 μm – 6.58 μm, depending on the value of *k*_1,*FR*_. After another six months, the outer diameter of the halo reaches 10.7 μm-11.4 μm.

**Fig. 4.**
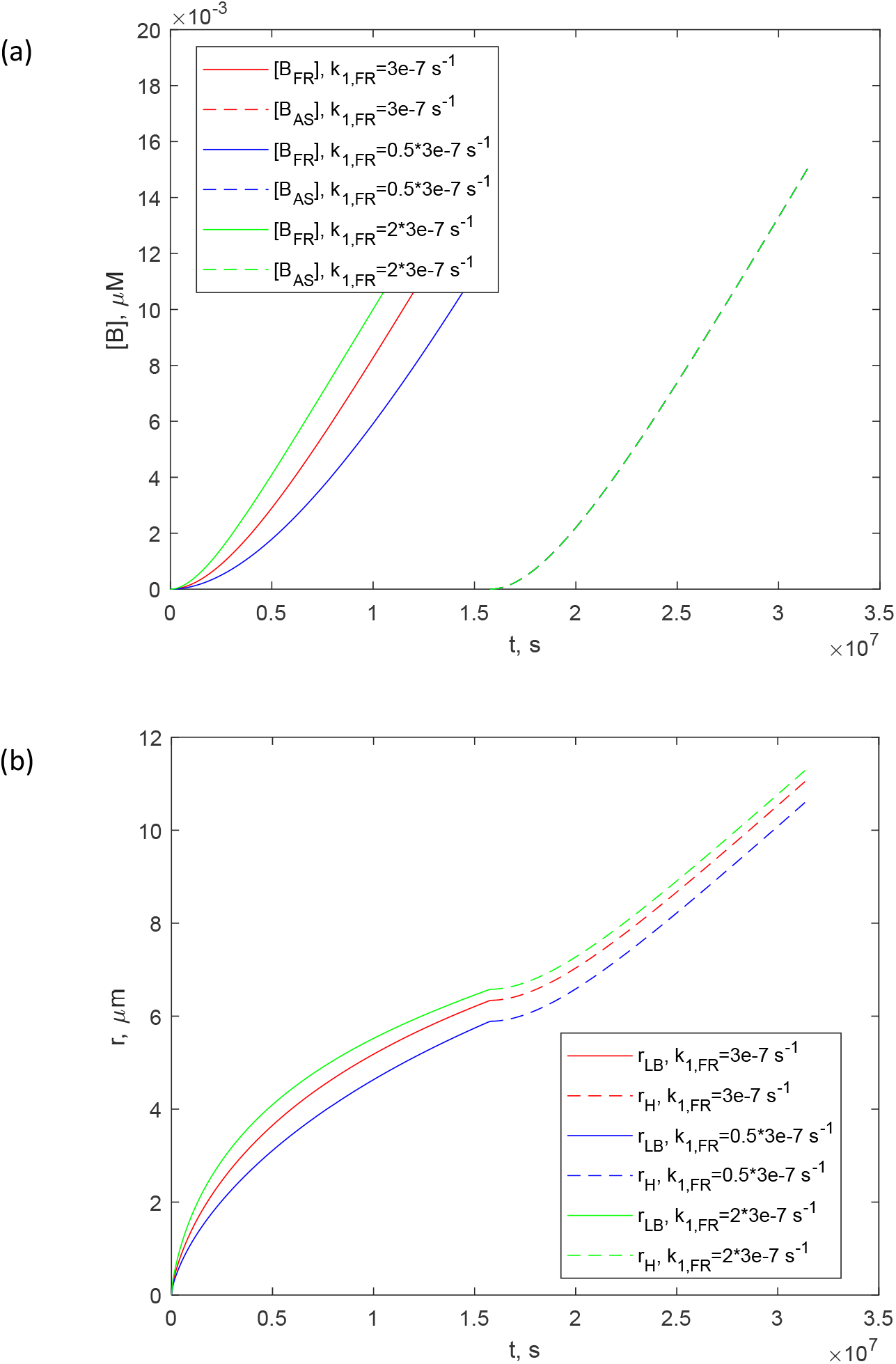
(a) Molar concentration of aggregated lipid membrane fragments in the core of the LB, [*B_FR_*], and molar concentration of α-syn fibrils in the halo of the LB, [*B_AS_*], versus time. (b) Radius of the core of the LB, *r_LB_*, and the outer radius of the halo of the LB versus time, *r_H_*, for three values of *k*_1,*FR*_. *q_FR_* = 5×10 ^-27^ mol s^-1^, *q_AS_* = 5×10^-27^ mol s^-1^, *k*_2,*FR*_ = 2×10^-6^ μM^-1^ s^-1^, *k*_1,*AS*_ = 3×10^-7^ s^-1^, *k*_2,*AS*_ = 2×10^-6^ μM^-1^ s^-1^.

An increase in the rate constant for the second pseudoelementary step of the F-W model describing autocatalytic attachment of lipid membrane fragments to the LB core, *k*_2,*FR*_, increases the rate of membrane fragment aggregation in the LB core (Fig. 5a). This results in faster growth of the core (Fig. 5b). It should be noted that for given parameter values the solution is less sensitive to *k*_2,*FR*_, and to produce visibly distinct curves in Fig. 5 we had to increase the value of *k*_2,*FR*_ by a factor of 10^3^. After six months of growth, the diameter of the LB core reaches 6.30 μm – 6.81 μm, depending on the value of *k*_2,*FR*_. After another six months, the outer diameter of the halo reaches 11.1 μm-11.6 μm.

**Fig. 5.**
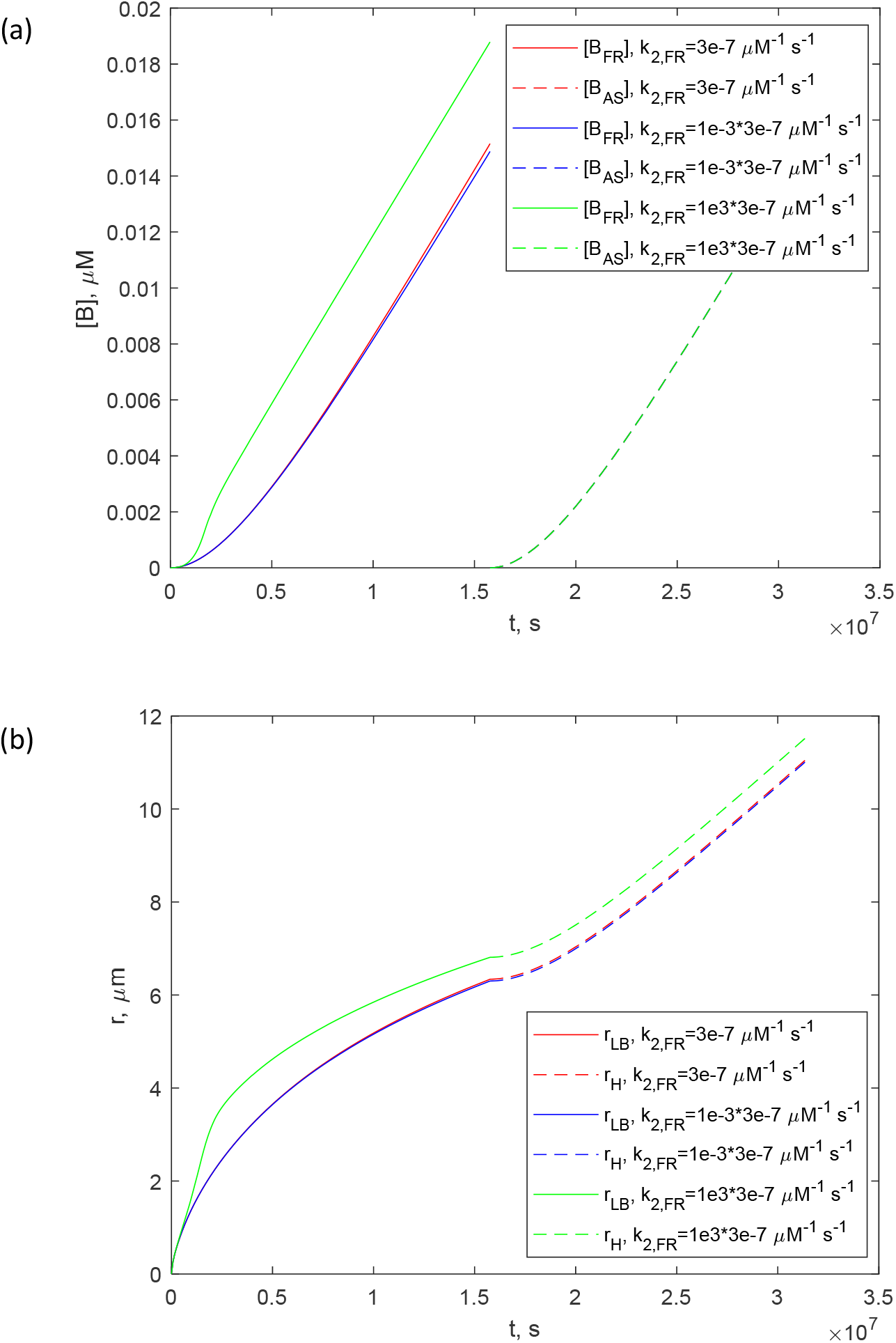
(a) Molar concentration of aggregated lipid membrane fragments in the core of the LB, [*B_FR_*], and molar concentration of α-syn fibrils in the halo of the LB, [*B_AS_*], versus time. (b) Radius of the core of the LB, *r_LB_*, and the outer radius of the halo of the LB versus time, *r_H_*, for three values of *k*_2,*FR*_. *q_FR_* = 5×10^-27^ mol s^-1^, *q_AS_* = 5×10^-27^ mol s^-1^, *k*_1,*FR*_ = 3×10^-7^ s^-1^, *k*_1,*AS*_ = 3×10^-7^ s^-1^, *k*_2,*AS*_ = 2×10^-6^ μM^-1^ s^-1^.

An increase in the rate constant for the first pseudoelementary step of the F-W model describing nucleation of α-syn aggregates, *k*_1,*AS*_, leads to an increase of the nucleation rate of α-syn aggregates (Fig. 6a). This leads to faster growth of the LB halo (Fig. 6b). After six months of growth, the diameter of the LB core reaches 6.34 μm. After another six months, the outer diameter of the halo reaches 10.2 μm-11.7 μm, depending on the value of *k*_1,*AS*_.

**Fig. 6.**
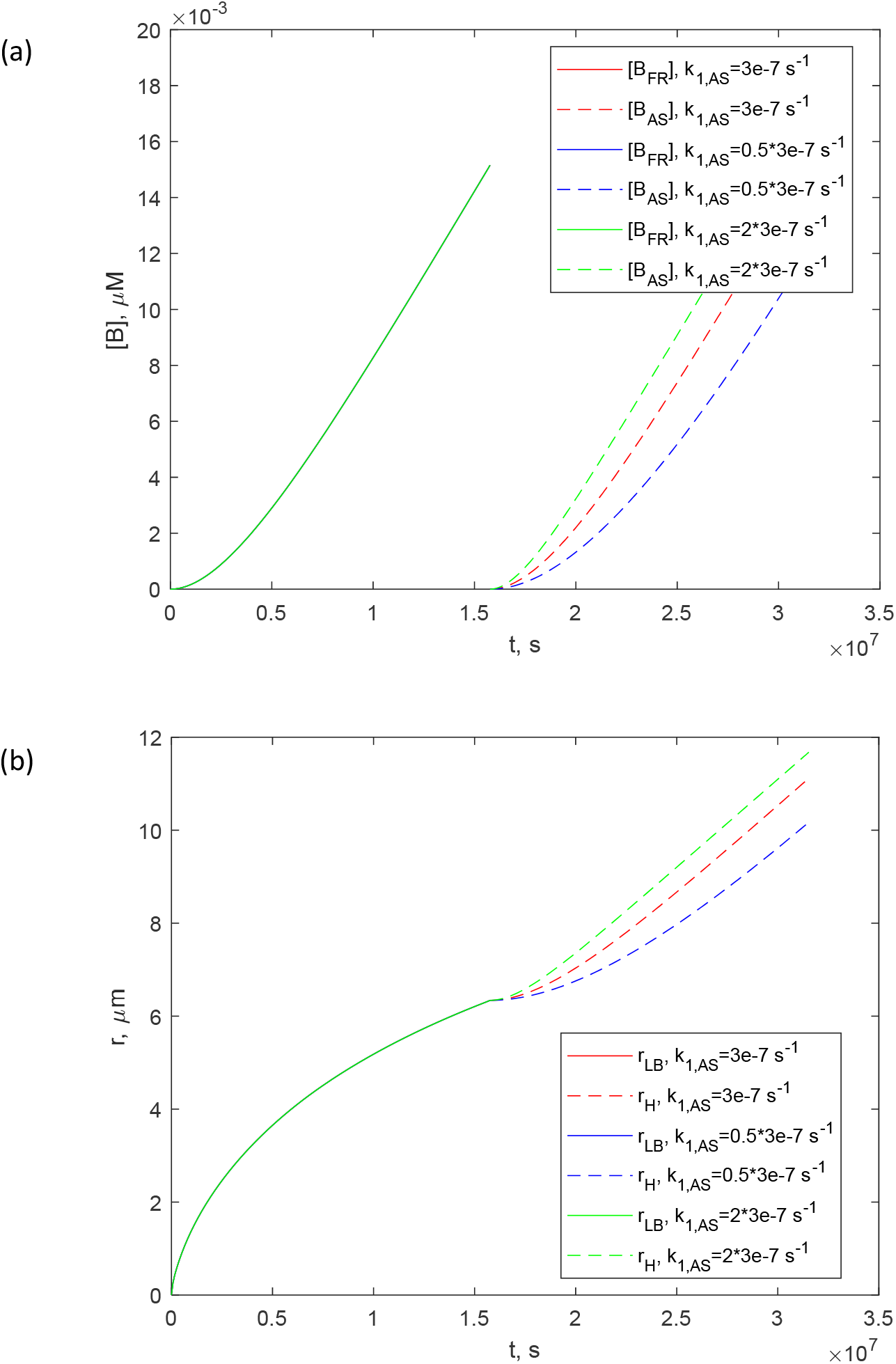
(a) Molar concentration of aggregated lipid membrane fragments in the core of the LB, [*B_FR_*], and molar concentration of α-syn fibrils in the halo of the LB, [*B_AS_*], versus time. (b) Radius of the core of the LB, *r_LB_*, and the outer radius of the halo of the LB versus time, *r_H_*, for three values of *k*_1,*AS*_. *q_FR_* = 5×10^-27^ mol s^-1^, *q_AS_* = 5×10^-27^ mol s^-1^, *k*_1,*FR*_ = 3× 10^-7^ s^-1^, *k*_2,*FR*_ = 2× 10^-6^ μM^-1^ s^-1^, *k*_2,*AS*_ = 2× 10^-6^ μM^-1^ s^-1^.

The increase of the rate constant for the second pseudoelementary step of the F-W model describing autocatalytic growth of α-syn fibrils due to attachment of α-syn monomers in the LB halo, *k*_2,*AS*_, leads to an increase of autocatalytic growth of α-syn fibrils (Fig. 7a). This leads to faster growth of the LB halo (Fig. 7b). As in Fig. 5, to produce distinctly different curves in Fig. 7 the value of *k*_2,*AS*_ had to be increased by a factor of 10^3^. After six months of growth, the diameter of the LB core reaches 6.34 μm. After another six months, the outer diameter of the halo reaches 11.0 μm-12.3 μm, depending on the value of *k*_2,*AS*_.

**Fig. 7.**
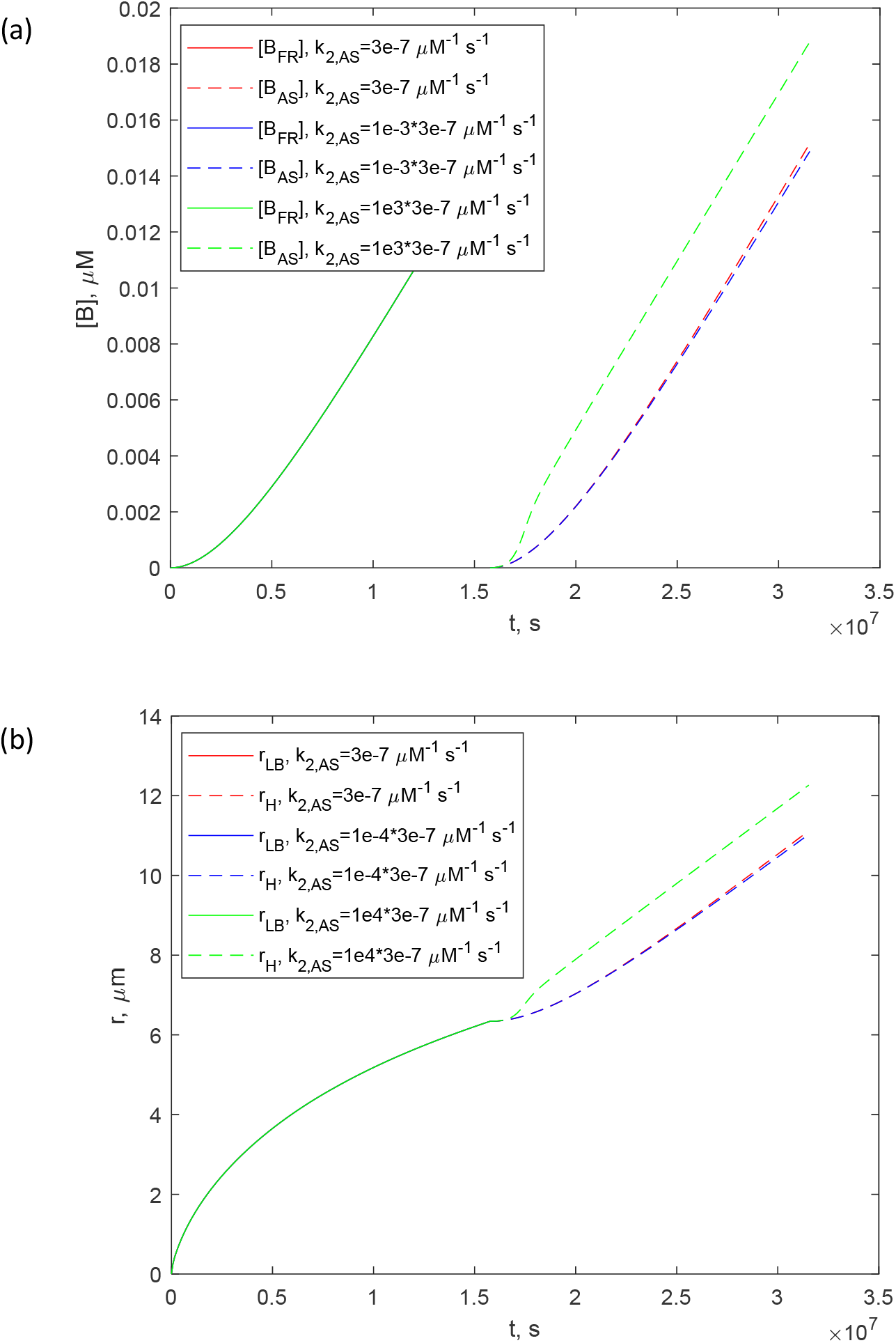
(a) Molar concentration of aggregated lipid membrane fragments in the core of the LB, [*B_FR_*], and molar concentration of α-syn fibrils in the halo of the LB, [*B_AS_*], versus time. (b) Radius of the core of the LB, *r_LB_*, and the outer radius of the halo of the LB versus time, *r_H_*, for three values of *k*_2,*AS*_. *q_FR_* = 5×10^-27^ mol s^-1^, *q_AS_* = 5×10^-27^ mol s^-1^, *k*_1,*FR*_ = 3×10^-7^ s^-1^, *k*_2,*FR*_ = 2× 10^-6^ μM^-1^ s^-1^, *k*_1,*AS*_ = 3×10^-7^ s^-1^.

### 3.2 Investigating the sensitivity of the size of the LB to model parameters

The relative sensitivity coefficients are reported in Table 4. An important parameter predicted by the model is the outer halo radius at the end of the simulation, *r_H,f_*. We investigated how *r_H,f_* depends on model parameters. The final radius of the LB is highly sensitive to the generation rates of membrane fragments and α-syn monomers. Also, *r_H,f_* is sensitive to the rate constants for the first pseudoelementary steps of the F-W model describing nucleation of membrane fragment aggregates and α-syn aggregates.

**Table 4.** Relative sensitivity of the outer halo radius at the end of the simulation (*t_H,f_* = 1 year), *r_H,f_*, to production rates of membrane fragments and α-syn monomers and kinetic constants characterizing aggregation rates of membrane fragments and α-syn monomers. Computations were performed using Eqs. (22) and (23) with (for example) Δ*q_FR_* = 10^-3^ *q_FR_*.

| $S_{q_{FR}}^{r_{H,f}}$ | $S_{q_{AS}}^{r_{H,f}}$ | $S_{k_{1,FR}}^{r_{H,f}}$ | $S_{k_{2,FR}}^{r_{H,f}}$ | $S_{k_{1,AS}}^{r_{H,f}}$ | $S_{k_{2,AS}}^{r_{H,f}}$ |
| --- | --- | --- | --- | --- | --- |
| 0.193 | 0.437 | 0.0428 | 0.00333 | 0.0968 | 0.00753 |

## 4. Discussion, limitations of the model, and future directions

The two main results obtained in this paper are the model of LB growth and an analytical solution of this model. The LB is assumed to consist of a core, which is composed of aggregated lipid membrane fragments, damaged organelles, lysosomes, and damaged mitochondria, and a halo, which is composed of radiating α-syn filaments.

The developed model is limited largely because of its minimalistic nature. Indeed, real LBs consist of more than 300 proteins (Shahmoradian *et al.*, 2019). More detailed mechanisms involved in the formation and growth of LBs need to be added in future research. Detailed biochemical simulations will require more experimental information including the values of kinetic constants, and future model development should be closely guided by experimentation. In addition, the F-W model does not track the size of polymers. A model capable of simulating polymers with different sizes would improve the value of the simulation. Future research should develop a physical model explaining why the growth of a lipid core is replaced by the growth of a halo that is composed of radiating filaments. There is likely a critical radius of the core when such a change becomes thermodynamically favorable. The question if LBs are neurotoxic or benign, simply serving as a sink for neurotoxic oligomers, remains one of the main questions in PD research. Future model development should seek to address this essential topic.

## Ethical Statement

The manuscript does not contain human or animal studies.

## Acknowledgment

IAK acknowledges the fellowship support of the Paul and Daisy Soros Fellowship for New Americans and the NIH/National Institute of Mental Health (NIMH) Ruth L. Kirchstein NRSA (F30 MH122076-01). AVK acknowledges the support of the National Science Foundation (award CBET-2042834) and the Alexander von Humboldt Foundation through the Humboldt Research Award.

